# Influential parameters for the analysis of intracellular parasite metabolomics

**DOI:** 10.1101/190421

**Authors:** Maureen A. Carey, Vincent Covelli, Audrey Brown, Gregory L. Medlock, Mareike Haaren, Jessica G. Cooper, Jason A. Papin, Jennifer L. Guler

## Abstract

Metabolomics is increasingly popular for the study of many pathogens. For the malaria parasite, *Plasmodium falciparum*, both targeted and untargeted metabolite detection has improved our understanding of pathogenesis, host-parasite interactions, and antimalarial drug treatment and resistance. However, purification and analysis procedures for performing metabolomics on intracellular pathogens have not been explored. Here, we investigate the impact of host contamination on the metabolome when preparing samples using standard methods. We purified *in vitro* grown ring stage intra-erythrocytic *P. falciparum* parasites for untargeted metabolomics studies; the small size of this developmental stage amplifies the challenges associated with metabolomics studies as the ratio between host and parasite biomass is maximized. Following metabolite identification and data preprocessing, we investigated whether host contributions could be corrected post hoc using various normalization approaches (including double stranded DNA, total protein, or parasite number). We conclude that normalization parameters have large effects on differential abundance analysis and recommend the thoughtful selection of these parameters. However, normalization does not remove the contribution from the parasite’s extracellular environment (culture media and host erythrocyte). In fact, we found that extra-parasite material is as influential on the metabolome as treatment with a potent antimalarial drug with known metabolic effects (artemisinin). Because of this influence, we could not detect significant changes associated with drug treatment. Instead, we identified metabolites predictive of host and media contamination that can be used to assess sample purification. Our findings provide a basis for development of improved experimental and analytical methods for future metabolomics studies of intracellular organisms.

## IMPORTANCE

Molecular characterization of pathogens, such as the malaria parasite, can lead to improved biological understanding and novel treatment strategies. However, the distinctive biology of the *Plasmodium* parasite, such as its repetitive genome and requirement of growth within a host cell, hinders progress towards these goals. Untargeted metabolomics is a promising approach to learn about pathogen biology. By measuring many small molecules in the parasite at once, we gain a better understanding of important pathways that contribute to the parasite’s response to perturbations like drug treatment. Although increasingly popular, approaches for intracellular parasite metabolomics and subsequent analysis are not well explored. The findings presented in this study emphasize the critical need for improvements in these areas to limit misinterpretation due to host metabolites and standardize biological interpretation. Such improvements will aid both basic biological investigations and clinical efforts to understand important pathogens.

## INTRODUCTION

Malaria continues to be responsible for hundreds of thousands of deaths annually, most of which result from infection with the protozoan parasite *Plasmodium falciparum* (1). Characterization of the biology of this important pathogen can lead to improved treatment strategies. ‘Omics approaches, such as genomics, transcriptomics, and proteomics, are widely used but the limited annotation of the parasite’s genome makes these data sets challenging to interpret. One way to alleviate this lack of functional knowledge is to use network-based modeling to contextualize noisy or sparse data and facilitate the interpretation of complex data (2). Additionally, the measurement of direct mediators of the phenotype, such as signaling and biosynthetic metabolites, can improve the ability to characterize phenotypes mediated by proteins that are not yet annotated into the genome. For this reason, metabolomics is becoming increasingly popular in studies of intra-erythrocytic stages of *P. falciparum* (3-12). These studies have improved our understanding of malaria pathogenesis (13), strain-specific phenotypes (11), and host-parasite interactions (9). Recent studies have successfully identified metabolic signatures that correlate well with biological function, such as time-and dose-dependent response to antimalarial treatment (3, 5) and resistance-conferring mutations (12).

Previous studies on *P. falciparum* have been confined to the larger, late intra-erythrocytic stage parasites. This is mainly due to available purification approaches; for example, magnetic purification specifically enriches late stage parasites that contain paramagnetic hemozoin while excluding early ring stages and uninfected host cells (14). Accordingly, the study of the smaller, early ring stage parasite is more challenging due to an inability to isolate adequate amounts of parasite material from host material (15). However, specific functionality can only be observed in the early parasite stages *(i.e.* artemisinin resistance) and metabolic details would greatly advance our understanding of such phenotypes.

There are distinct challenges that need to be considered when performing metabolomic studies in obligate intracellular pathogens, like *P. falciparum;* chief among these are (1) acquiring adequate material and (2) the potential for contamination from host cells. Due to inefficient purification methods, samples typically have few parasites and yet abundant host erythrocyte material. Uninfected host cells are often >10 times more prevalent than *P. falciparum-infected* host cells in laboratory culture and clinical infections, and the host erythrocyte contains up to 10-fold more cellular material (16, 17).

In this study, we sought to define critical parameters that can be used to overcome these challenges and facilitate the collection of high quality metabolomics data. We chose to investigate an extreme case, metabolically perturbing early ring-stage *P. falciparum* parasites, to determine if extensive extra-parasite contamination present after commonly used isolation methods can be removed analytically. We show that both the choice of analytic parameters (in particular, the normalization approach) and extra-parasite contamination heavily influence the interpretation of metabolic changes. However, even appropriate normalization fails to remove environmental noise completely. Contamination from the media and host cells is as influential on metabolome as sample treatment. Thus, we propose that the combination of improved purification and analytic parameters will generate more accurate measures of the metabolome, increasing the utility of untargeted metabolomics to investigate intracellular parasite biology.

## RESULTS

### Metabolomics

We conducted metabolomics on early ring stage (0-3 hour) *Plasmodium falciparum* parasites lysed from host erythrocytes. Two parasite clones were grown in matched conditions, lysed and washed from the host cell, and analyzed via Ultrahigh Performance Liquid Chromatography- coupled with mass spectrometry (Fig. 1A). Prior to isolation, each clone (either a drug sensitive and resistant line) was either untreated or treated with 700nM dihydroartemisinin (for 6 hours), generating four sample groups with matched blood batches, media, and purification approach (Fig. 1B). Dihydroartemisinin, the active component of the antimalarial artemisinin, is a known metabolic disruptor (3, 5, 18). Both sensitive and resistant parasites are known to enter a unique metabolic state, called dormancy, following treatment. Dormancy is characterized by reduced metabolic activity (19-21), and, thus, treated ring-stage parasites should have a distinct metabolome when compared to untreated parasites.

**Figure 1.**
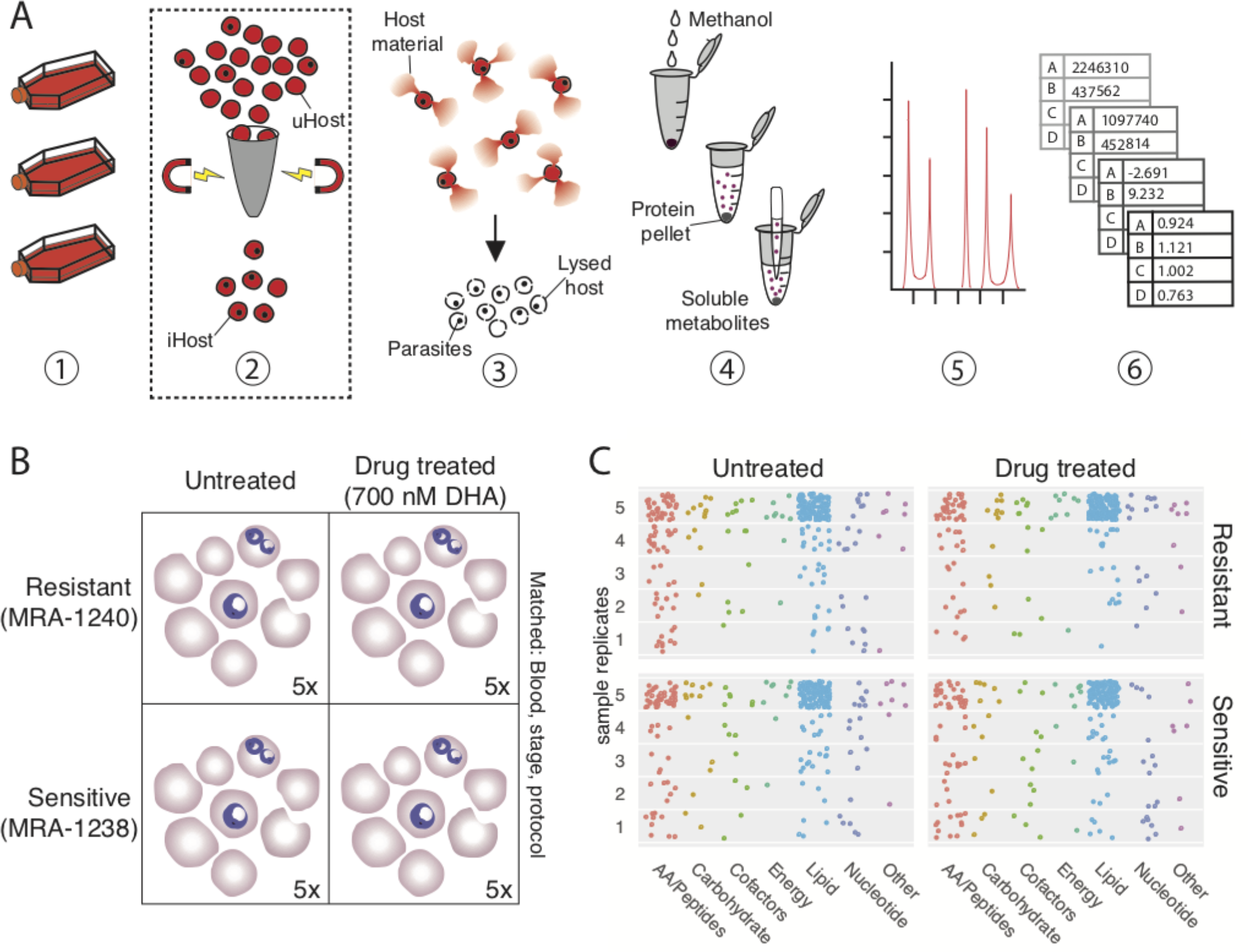
Metabolomics pipeline and metabolite identification. **A: Metabolomics purification and analysis pipeline**. 1) Laboratory-adapted *P. falciparum* clones are cultured in host erythrocytes. Parasite count is collected at this step. 2) If enriching for late stage parasites, cultures are passed through a magnetic column to retain paramagnetic late stage-infected erythrocytes. Note: this was not done for this study. iHost; infected host erythrocytes, uHost; uninfected host erythrocytes. 3) Host erythrocytes are lysed using saponin, but parasites remain intact. Samples are washed to remove hemoglobin and other intracellular host material. Total protein is quantified at this step. 4) Soluble metabolites are extracted from precipitated protein using methanol and centrifugation. Double stranded DNA is quantified at this step. 5) Metabolites are separated via liquid chromatography and identified using mass spectroscopy. Metabolite spectra are compared to a library of authenticated standard metabolites for high-confidence identification. 6) Abundance data for each metabolite is normalized to an appropriate parameter (i.e. DNA content or parasite number), log transformed, centered to median, and scaled to variance, prior to employing statistical comparisons. **B: Experimental comparison**. All samples were grown in RPMI media supplemented with AlbuMAX and hypoxanthine and one of three blood batches. At the early ring stage (<3 hours post invasion), 10 samples were treated with dihydroartemisinin (DHA, 700nM) for 6 hours and 10 were protocol and condition matched (blood batch, media batch, and stage) without drug treatment (see Table S3). **C: Summary of identified metabolites**. Metabolites (each point) from various metabolic subgroups are not uniformly detected in all five replicates for any sample group. Consistency of measurement is indicated by the metabolite point placed in 1-5 replicates (y axis). The majority of metabolites detected were lipid species, as indicated by number of blue dots. A full list of identified metabolites is in supplemental data.

Mass spectrometry analysis of these samples detected 297 identifiable metabolites; 155 metabolites were detected in every sample. Samples contained between 182 and 267 metabolites. Detected metabolites represented 10 energy associated metabolites, 159 lipid species, 108 peptides and amino acids, 40 nucleotides, 28 cofactors, 20 carbohydrates, and 10 others (Table S1). Lipid species were most consistently detected in every sample (as measured by percentage of metabolite found in every sample) and amino acids were often unique to individual samples (Fig. 1C). Several metabolites were measured that have not previously been reported in *P. falciparum* including kynurenine (detected in 25% of samples), phenol red (phenolsulfonphthalein, in 95% of samples), and HEPES (4-(2-hydroxyethyl)-1-piperazineethane-sulfonic acid, in all samples; see Table S1).

### Host persistence

Despite implementing current best practices, including erythrocyte lysis and washing steps to remove parasites from their intracellular milieu (Fig. 1A, (3, 8, 12, 22)), parasite separation from the host is poor. Microscopy revealed that parasites lysed from host cells remain embedded in erythrocyte membranes and washes fail to isolate parasite material (Fig. 2A). Over 68% of parasites remain associated with host membrane (Table S2). This result emphasized that erythrocyte ‘ghosts’ (cell membranes with associated metabolites) remain abundant in the sample and heavily contribute to the metabolome. Thus, we sought analytic approaches to remove host contamination *post hoc*.

**Figure 2.**
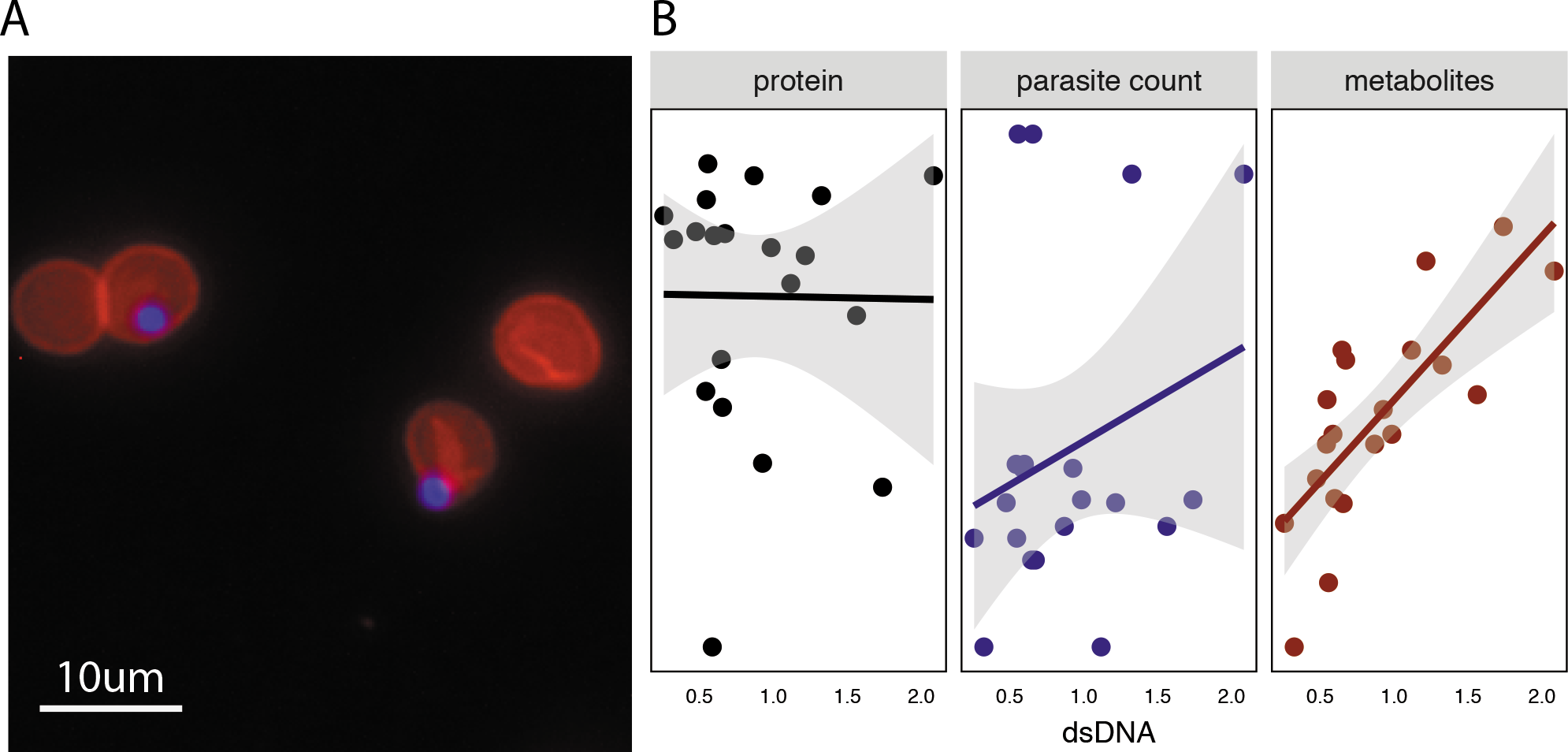
Host persistence is detected using multiple approaches. A: Visualization of parasites within erythrocyte ghosts. Fluorescent imaging (40X) reveals parasites (blue, DAPI) retained within erythrocyte ghosts (red, phycoerythrin conjugated CD235a antibody) following saponin treatment. Approximately 70% of parasite remain associated with host membranes (see Table S2). **B: Sample characteristics**. Samples were evaluated for double stranded DNA (dsDNA, quantified on x-axis), protein amount (black, quantified on y-axis in left panel), and parasite count (blue, quantified on y-axis in center panel) prior to analysis. Total number of metabolites detected per sample (red, y-axis) is significantly correlated with sample dsDNA quantification (p = 9.8*10^-5^, r^2^ = 0.76). Protein amount and parasite count are not significantly correlated with dsDNA. Fit line uses a linear model, and shaded region represents the standard error.

### Normalization

We first explored the use of normalization with three distinct approaches. Metabolomics preprocessing methods can influence results (23, 24) but the role of normalization has not been extensively explored, particularly in intracellular pathogens. Both host and parasite-derived metrics (double stranded DNA [dsDNA], protein, and parasite count) were evaluated in the experimental setup (Fig. 1A). Sample replicates contained 1.3-6.9 million parasites (Table S3). As expected, no two normalization metrics were correlated across samples (Fig. 2B; see supplementary information for code). Metabolite yield (as measured by number of identified metabolites) was only correlated with DNA abundance (p = 9.8*10^-5^, r^2^ = 0.76; Fig. 2B), indicating DNA abundance is best associated with total biomass.

Initially, we anticipated that dsDNA should come primarily from the parasite fraction, as host erythrocytes are anucleated and growth media does not contain any intact DNA; however, we found that host cells and AlbuMAX (a media component) do contribute to sample dsDNA (Fig. S1). Protein is likely derived from all three culture components as well: parasite, host erythrocyte, and media (via AlbuMAX supplementation). Although parasite count is a direct measure of the parasite fraction, this variable was collected several steps upstream of metabolome quantification (Fig. 1A) and may be suboptimal when compared to metrics collected later in the pipeline.

Accordingly, we normalized metabolomes to these parasite and host-derived metrics to determine if normalization reduces extra-parasite noise to reveal parasite metabolomes. Normalization of metabolite levels can be calculated by a variety of methods (Table 1, Fig. 3), all aiming to enhance interpretation of results by controlling for technical or non-biological variation. To normalize, we divide the abundance of each metabolite in a sample by its corresponding sample variable to control for sample-to-sample variation (Fig. 3). As illustrated in Figure 3, normalization can significantly affect interpretation of results and should be selected carefully based on experimental design and knowledge of samples.

**Figure 3.**
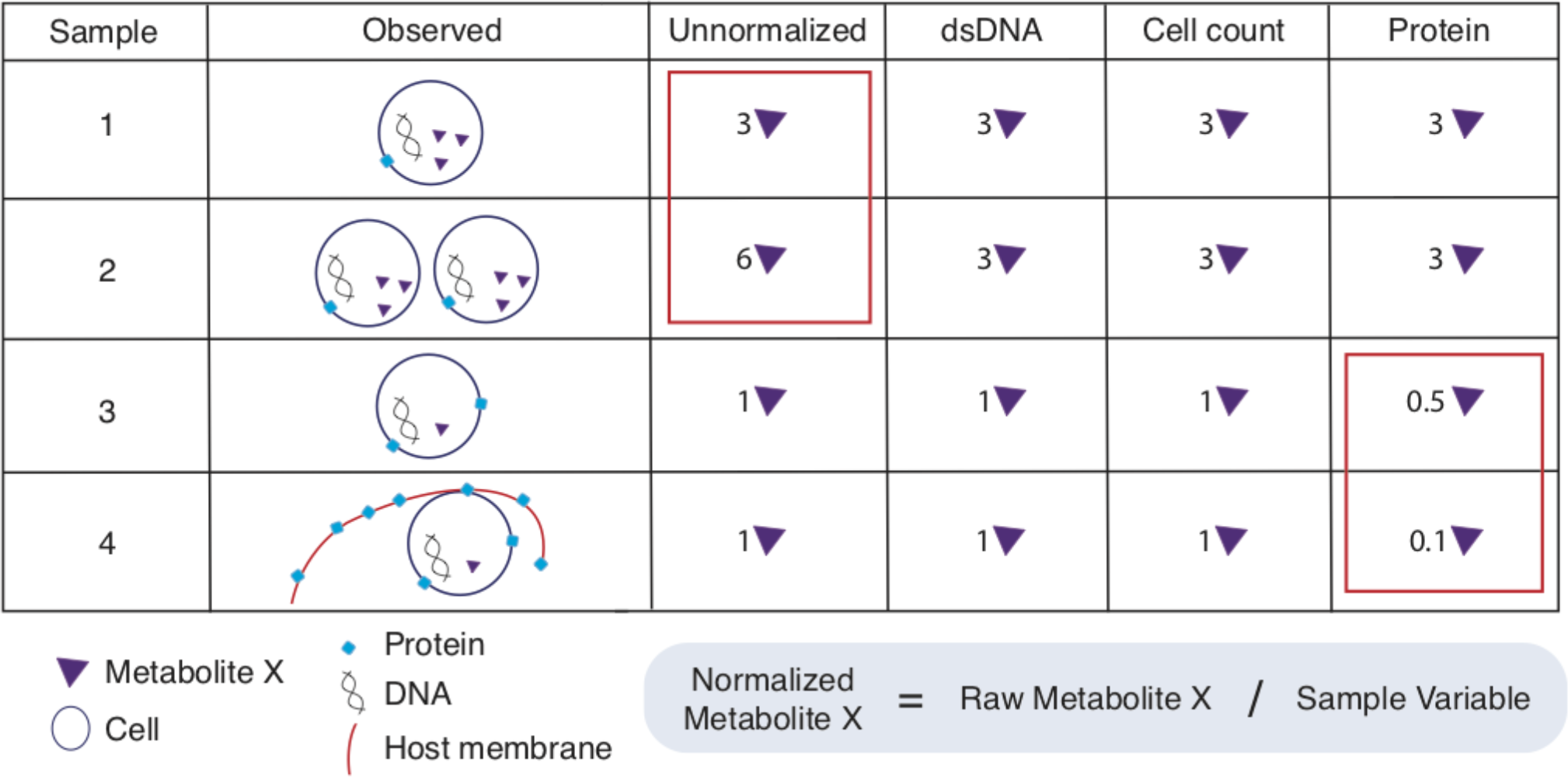
Normalization approaches impact final metabolite abundance. Normalization controls for sample-to-sample variation. Normalization requires sample metabolite abundance to be divided by quantified normalization factor, the sample variable (equation in blue box, normalization factors above box). Example results shown in table indicate abundance of metabolite X given several different sample metrics for normalization. For example, identical samples with difference cell count (sample 1 and 2) reveal the importance of normalization; without it, the identical samples have a 2-fold difference in metabolite X. Identical parasite samples 3 and 4 also have a nearly 2-fold difference in metabolite abundance when normalizing to protein, due to host bias on protein measure.

**Table 1.** Parameters in metabolomics analysis of intracellular parasites, including *Plasmodium*. Note: most parameters do not have strict recommendations, as they are dependent on experimental design. Bolded text indicates methods that were employed and evaluated during this study.

| Parameter | Options | Factors to consider |
| --- | --- | --- |
| Growth conditions | **Ring stage** | **-Limited biomass (1-2µm, Figs. 1A and B), haploid genome**<br>**-Few enrichment options** |
|  | Late stage | -Larger in size (3-10µm), polyploid genome<br>-Can use magnetic enrichment (Fig. 1A) |
|  | Mixed stages | -Consider effects of stage variation on data |
|  | Media batches | -Relevant if using serum-based media formulations |
|  | **Blood batches** | **-Must be recorded and matched within comparisons (Table S3)**<br>**-Useful to assess host contamination levels (Fig. 5 & 6)** |
| Additional controls | Uninfected erythrocytes | -Use to identify host metabolites<br>-Used in addition to normalization |
| Enrichment methods | **Saponin, other lytic reagents** | **-Compatible with all stages (Fig. 1A)**<br>**-Parasites remain in erythrocyte ghosts (Fig. 2A)**<br>**(Need improved methods that isolate parasite from host cell)** |
|  | Magnetic purification | -Increases parasite to host ratio (Fig. 1A) |
| Metabolite Detection | NMR | -Limited metabolite detection but higher confidence |
|  | **Mass Spectrometry** | **-Industry standard for broad detection** |
|  | Radio labeling | -Targeted approach with high confidence |
|  | Single metabolite assays | -High-confidence, targeted approach with low throughput |
| Pre-analysis normalization | Cell number normalization | -Can be combined with any post-analysis normalization but requires sample manipulation |
| Post-analysis normalization | **Parasite-derived parameters** | **-i.e. parasite number**<br>**-Selection requires knowledge of experimental design** |
|  | **Parameters with mixed derivation (host, parasite)** | **-i.e. protein, DNA**<br>**-Can fail to remove undesired noise (Fig. 2 & 4)** |
|  | Internal standards | -Dependent on metabolomics facilities |
| Centering | Mean | -Standard centering |
|  | **Median** | **-Less sensitive to outliers** |
|  | Other | -See (24) for summary of alternative approaches |
| Scaling | **Within group SD** | **-Requires no additional samples** |
|  | Z-scoring | -Requires control samples (*i.e.* untreated or uninfected erythrocytes) |
| Statistical analysis | **Univariate** | **-Requires multiple comparison corrections** |
|  | **Multivariate** | **-Reveals group differences based on multiple variables** |
|  | **Machine learning (e.g. Random Forest)** | **-Classification is more stringent than univariate tests, but can identify nonlinear effects** |

Because the effect of normalization has not been explored in intracellular parasites, we normalized to parasite number (parasite-derived), dsDNA (parasite, media, and host derived), and total protein amount (parasite, media, and host derived), and then performed principal component analysis with all sample metabolomes (Fig. 4A-D). Each normalization method yields distinct principle component structures, yet none clearly separate the four sample groups (as measured by clustering of the sample groups by PerMANOVA, p-values provided in figure under normalization heading). However, with DNA normalization, we are able to separate drug treated parasites from untreated parasites or clonal groups (Fig. 4B); with parasite number normalization, we can distinguish clonal groups (Fig. 4D).

**Figure 4.**
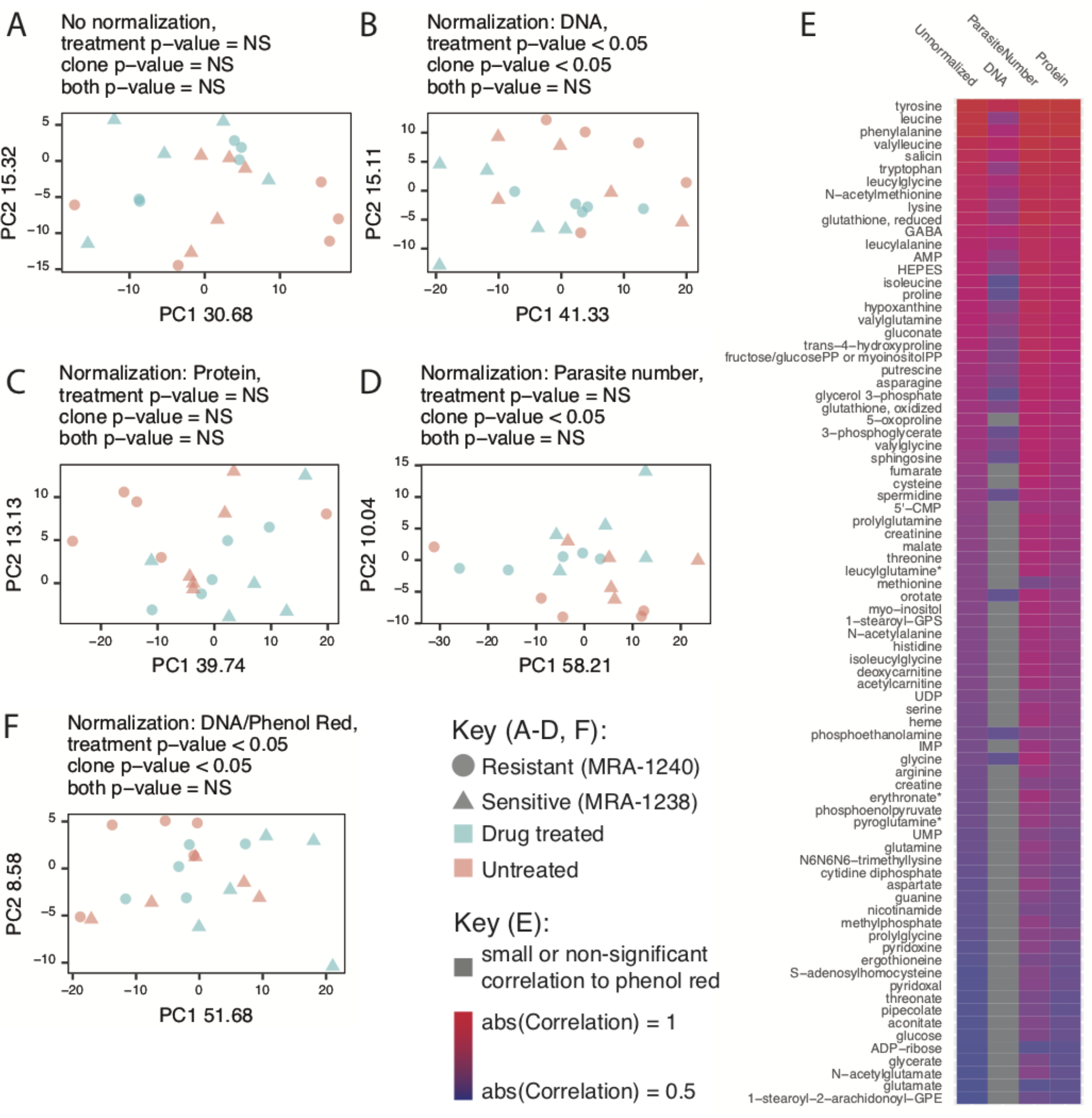
Metabolomes are dependent on normalization approach and influenced by extra-parasite contamination. A-D: Normalization affects metabolome similarity. Principle component (PC) analysis was performed prior to normalization (**A**), as well as after using three different normalization methods (DNA (24), total protein (24), and parasite count [**D**]). PerMANOVA significance is listed for each grouping. **E: Metabolites associated with media components.** The raw abundance of 82 metabolites is correlated with phenol red (unnormalized column). These associations are not removed with parasite number and protein normalization. DNA normalization best removes associations with media components (increase in number of grey or insignificant correlations); only 39% of correlations remain. **F: Removal of media-associated metabolites.** PCA of DNA normalized samples with phenol red correlated metabolites removed from the dataset yields no improvement in sample clustering.

Consistent with the lack of distinct separation, univariate statistical analysis revealed no differentially abundant metabolites between the four groups (see supplemental information for code). When normalization is employed, metabolome differences between groups are highly dependent on approach; the top differentially abundant metabolites are normalization method-dependent (*data not shown*; see supplemental information for code). These findings emphasize that biological interpretation can change significantly depending on the chosen analytic parameters, and, thus, the selected normalization metric is a critical parameter and must be shared for analytic reproducibility.

### Data filtering

We next examined extra-parasite metabolites in our data set to estimate, and potentially correct for, the level of host contamination. Because there are no unique metabolites associated with the host, we explored media-specific metabolites, specifically phenol red and HEPES. Both phenol red and HEPES are components of the growth medium and should not be utilized by host or parasite. These metabolites are routinely excluded from metabolomics analysis for this reason.

Interestingly, the abundances of 82 and 76 metabolites (out of 298 total metabolites) are correlated with phenol red or HEPES, respectively (Fig. 4E and *data not shown*); the abundances of 59 metabolites are correlated with both compounds. Many of these metabolites remain correlated with the media components even after normalization (phenol red shown in Fig. 4E, >39%). Because both of these media metabolites appear to increase in abundance in drug treated samples (nonsignificant trend, *data not shown*), we argue that this extra-parasitic fraction will influence the interpretation of drug treatment. If we remove these media components and associated metabolites from our analysis and focus on the remaining 216 metabolites, sample separation into the four treatment groups surprisingly does not improve over DNA normalization alone (Fig. 3F). Thus, both *post hoc* normalization and data filtering are likely insufficient to remove the effect of extra-parasite contamination, especially in low-powered studies.

### Machine learning

We next used machine learning to attempt to separate the extra-parasite-associated metabolome from the parasite metabolome. Here, we leveraged the multiple blood batches used in parasite culture (Fig. 1A). Our four sample groups were grown in three different blood batches (Table S3). Univariate statistical analysis revealed only one metabolite with differential abundance among blood batches (1-arachidonoyl-GPE; see supplemental information for code).

To further explore the host contribution to the metabolome, we built Random Forest classifiers to predict blood batch or drug treatment (Fig. 5). Random Forest analysis is an internally validated machine learning approach, used here to classify samples into groups based on their metabolome (Fig. 5A) and to identify individual variables that are important for prediction accuracy (Fig. 5B).

**Figure 5.**
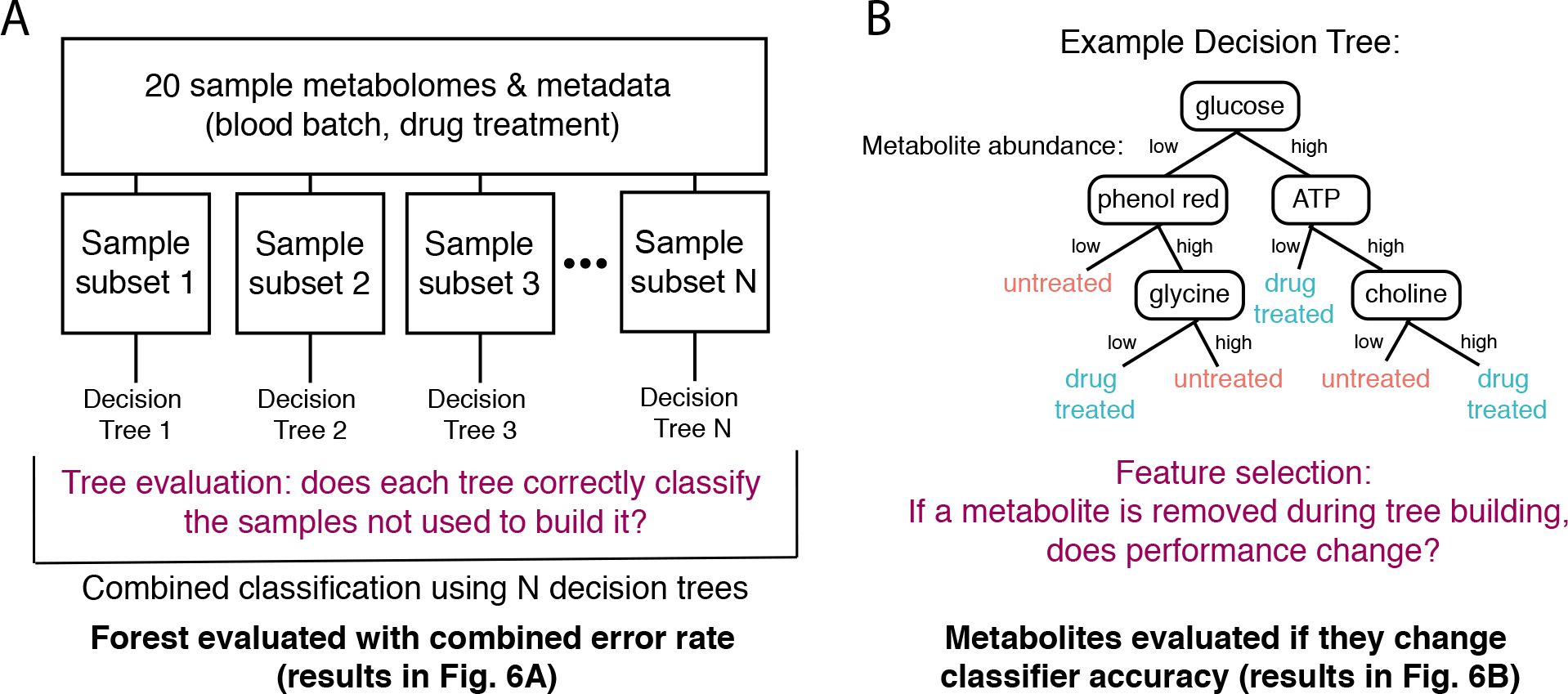
Random Forest Analysis. A: Building a random forest classifier. Samples are randomly subsetted into training and test datasets; from the training subsets, decision trees are built to separate samples into groups (see panel **B**). Trees are evaluated by testing classification performance on remaining samples from test datasets. See methods for more detail on analysis. **B: Evaluating metabolite importance.** Metabolite importance is calculated by determining the effect on classifier performance when the metabolite is removed from the dataset. See methods for further detail.

We first built a classifier to predict blood batch across all samples. Ninety-five metabolites (of 298) improved classifier accuracy in predicting blood batch (see supplemental information for code). Many of these metabolites are correlated in abundance with the media components explored (Fig. 4E), including CDP-ethanolamine, AMP, ADP-ribose, and aspartate which are among the top ten most influential metabolites in this classifier. The remaining metabolites had no effect on classifier performance or worsened its predictive ability, indicating they are not associated with blood batch due to high variability or association with other features that differentiate samples. This classifier built from DNA-normalized metabolomes predicted blood batch with a 30% error rate (Fig. 6A).

**Figure 6.**
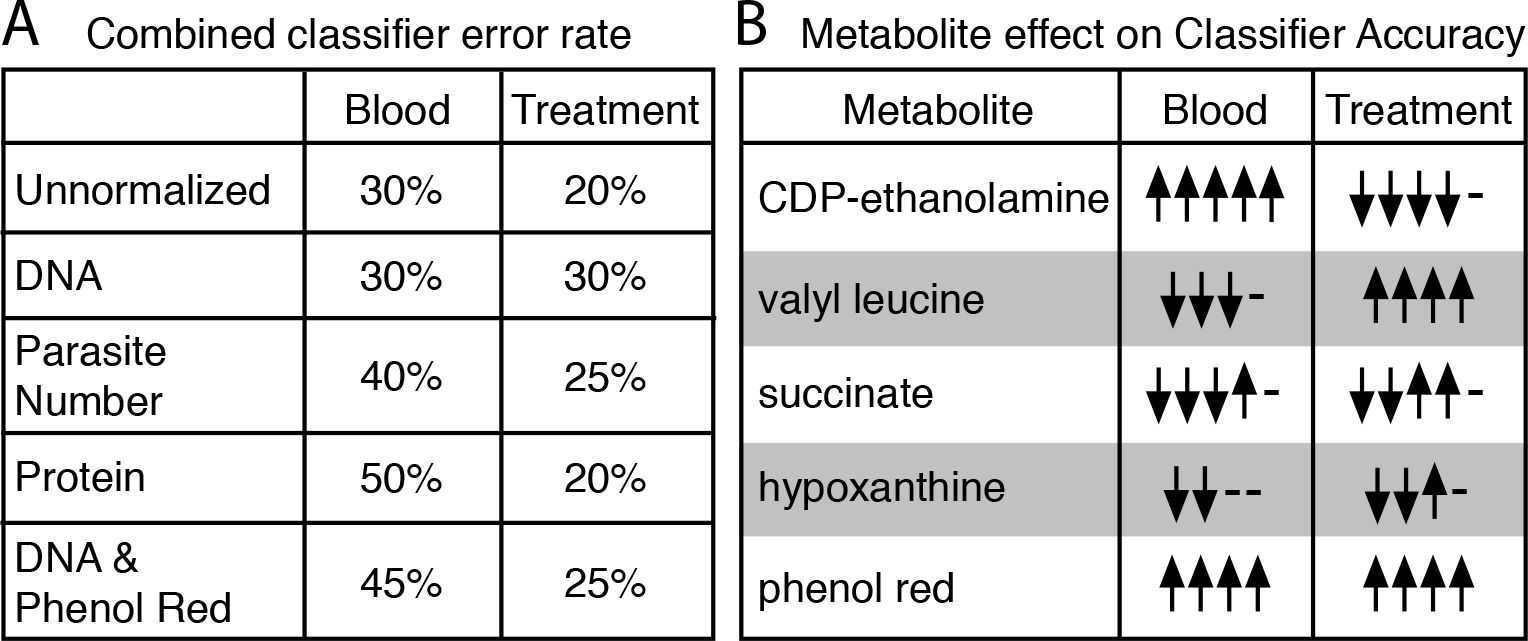
Blood batch and antimalarial treatment influence metabolomes. **A: Classifier performance**. Classifiers were built to predict blood batch or treatment condition using the metabolomics data with or without 4 normalization approaches. Classifier error rate varies with normalization approach. **B: Normalization method determines important metabolites.** A sample of five metabolites associated with improved or worsened classifier accuracy are shown. These metabolites are shown due to their importance in classifier performance and interesting behavior across classifiers. Up arrows indicate the metabolite improves classifier accuracy in one classifier and down arrows indicate they worsen accuracy in one classifier (arrows represent normalization approaches from panel A); if the metabolite does not improve or worsen accuracy, a dash is shown. Contradictory results (both up and down arrows for one metabolite) indicate that the normalization method changes metabolite importance. Note, valyl leucine, hypoxanthine, and phenol red were removed upon phenol red filtering and, therefore, are only present in 4 classifiers, as indicated by the four arrows and dashes.

To determine if blood batch is as influential on metabolome as a potent antimalarial drug treatment, we built a similar classifier to predict artemisinin treatment. Parasites were classified into two treatment conditions with a 30% class error rate using DNA-normalized metabolomes (Fig. 6A). One hundred and eighteen metabolites (of 298) improved the accuracy of this classification, including media-correlated metabolites like pipecolate, several dipeptides, and phenol red (see supplemental information for code).

Our classifier performance (Fig. 6A) is relatively poor due to small sample size and indicates only a subset of the measured metabolomes was predictive of blood batch or drug treatment. Classifiers built from data under alternative normalization approaches were comparable in performance. Phenol red correction reduced blood batch classifier performance more than the treatment classifier; this result is not surprising because both media components and host cell are extra-parasitic. Thus, by removing media contamination, we may also be removing host contamination and data associated with blood batch. However, phenol red is associated with both blood batch and treatment classifier accuracy (Fig. 6B); this result supports the need to remove extra-parasitic metabolites during sample preparation as they can skew meaningful biological interpretation.

Interestingly, if the classifier was built using a different normalization approach, the set of metabolites that most contributed to accuracy changed (representative examples in Fig. 6B, code for full analysis in supplementary information). Although some metabolites (like CDP-ethanolamine or valyl leucine) are consistently associated with blood or treatment classifier accuracy, respectively, some metabolites have contradictory results depending on the data normalization approach (like succinate and hypoxanthine; Fig. 6B). Thus, sample metabolome can classify both blood batch and sample group, indicating that sample treatment and blood batch influence the metabolome, and this is normalization-approach dependent.

## DISCUSSION

The lifestyle of intracellular parasites presents challenges to implementing traditional metabolomics protocols, predominately due to host metabolite contamination and limited amounts of parasite material. These challenges are exacerbated when studying early parasite stages (like the *Plasmodium* ring stage here), when the parasite is smallest. In our study, we conducted a detailed assessment of the impact of extra-parasite contamination and investigated analytic approaches to improve metabolome interpretation. We recommend improved discussion of normalization methods in the metabolomics field, especially for intracellular parasites, as normalization significantly effects the interpretation of a dataset. Additionally, we propose several analytic approaches to explore the effect of host contamination.

### Metabolome interpretation is normalization-approach dependent

Normalization limits non-biological variation and is absolutely essential for biological interpretation (Fig. 3). Normalization factors can be calculated using a variety of methods and is implemented either before or after metabolite quantification and identification (described as pre-analysis or post-analysis, Table 1 (25, 26)). Often, pre-analysis normalization is conducted by isolating the same number of cells for analysis (27) but this is not typically used in the study of *P. falciparum* as generating adequate biomass can be challenging (3, 5, 12). Furthermore, the use of inaccurate quantification methods may negate this step by introducing more variability. Post-analysis normalization methods are also routinely used; these include the use of internal standards (26, 28), corrections for protein amount (often used for supernatant or cell-free metabolomics (29)), DNA content (an approach validated in mammalian cells (30) and applied to bacterial cells (31)), or cell number (typically used for bacterial populations (32)).

To our knowledge, normalization has not been described in detail in previous metabolomics studies of *P. falciparum*, perhaps due to the technical challenges that we explore here. We evaluated three post-analysis normalization approaches: protein, double stranded DNA, and parasite number (Fig. 3 and 4A-D). Overall, we conclude normalization significantly affects the interpretation of results (Fig. 4 and 6). Normalization approach influences the metabolites with greatest differential abundance *(data not shown* because they did not reach significance) and the metabolites predictive of sample group shift with data normalization (Fig. 6).

In the present studies, only parasite count is entirely parasite-derived. The extracellular environment (including media components and host erythrocyte) likely contributes heavily to protein abundance. Accordingly, parasite count and protein abundance are not correlated. We also show that the host cell contributes to dsDNA, despite lacking a nucleus (Fig. S1). Despite this finding, our analysis shows that dsDNA normalization of early ring stage metabolomes best distinguishes sample and treatment groups and removes media contamination (Fig. 4). Much variability still remains after this step; we did not identify any differentially abundant metabolites even though artemisinin has reported metabolic effects on late stage parasites (3, 5, 18) and dormancy induces metabolic shifts in ring stage parasites (19-21). Although dsDNA normalization is most effective in our dataset, it is not appropriate in all experimental cases; for example, this type of analysis would introduce variability when comparing groups of different parasite stages due to known genome copy number differences (33-35).

### Media and host contribute to the measured metabolome

We found that extra-parasite material contributed by host erythrocytes and media components can also heavily impact the metabolome. Many studies employ erythrocyte lysis prior to sample purification ((3, 8, 12, 22), see *Materials and Methods).* However, several novel results from our study show that this step does not eliminate the potential for host contamination.

First, lipid species were the major class of metabolites detected in our analysis (Fig. 1C) perhaps due to the abundance of erythrocyte membranes or ‘ghosts’ present in the preparations (Fig. 2A). Second, more than a quarter of the metabolome is correlated with the media components (phenol red, Fig. 4E, and HEPES, *data not shown*). These metabolites are neither produced or consumed by the parasite but likely remained associated with cells following *in vitro* culture in media that contains these metabolites (as well as high levels of other metabolites such as glutathione, hypoxanthine, glutamine, and many amino acids). Third, we measured metabolites not expected to be produced or consumed in *Plasmodium* (2). For example, kynurenine is present in erythrocytes, derived from the amino acid L-tryptophan (36, 37); it is not involved in *P. falciparum* metabolism (38). Lastly, the only differentially abundant metabolite in our entire analysis that reached significance was associated with blood batch (1-arachidonoyl-GPE). This metabolite has not been explored in erythrocyte or *Plasmodium* metabolism but can be explored as a potential marker of host contamination.

In fact, we were able to predict a set of metabolites that are most likely to be influenced or derived from the host erythrocyte by identifying the metabolites that are most predictive of blood batch (Fig. 5B and 6B, see figures in supplementary code for comprehensive list). Going forward, it may be possible to use this approach to use specific metabolite markers to (1) assess levels of host contamination and parasite sample purity and (2) control for host contamination during analysis.

### Future recommendations

In this study, we compiled evidence of host cell and media contamination in untargeted metabolomics studies of intracellular parasites. We showed that analytic approaches can improve the accuracy and interpretability of intracellular parasite metabolomes but, ultimately, better purification methods are needed to extract biological differences from samples.

A common approach used in the study of *P. falciparum* involves an uninfected erythrocyte control to adjust for the presence of host metabolites (7, 10, 22, 28, 39, 40). While this approach has been used in many studies, it is not always financially feasible to match each blood batch with an adequate number of replicates. We hypothesize that an uninfected erythrocyte control is not sufficient because it is used by removing metabolites identified in the uninfected host from the analysis of parasitized host, like our normalization with phenol red-associated metabolites removed (Fig. 4F). Thus, interpretation of data even with the use of this control remains challenging (*e.g.* (41)). Thus, we propose that similar to studies in *Leishmania* (42-44), normalization and discussion of the chosen normalization metrics should become standard during metabolomics analysis of intra-erythrocytic parasites. Another common analytic step involves the removal of extra-parasitic metabolites, such as phenol red, as they are considered to be noise from culture media. We propose that these metabolites should not be excluded from the dataset and subsequent analysis because they contain valuable information about experimental variation and could be used for quality control, as indicated by the frequent correlation between phenol red abundance and other metabolites (Fig. 4E).

Overall, the methodology and findings from the current study provide a basis for the improved use of *in vitro* metabolomics approaches for the future investigation of intracellular pathogens, including *P. falciparum.* We suggest a set of considerations and recommendations for enhancing the accuracy of parasite metabolomics (Table 1 and below). First, samples must be better purified away from host material. Purification could involve enrichment methods to increase parasitemia prior to lysis (reducing the ratio of uninfected host cells to parasites) or the direct removal of host material post lysis. Currently, only enrichment approaches for late stage malaria parasites exist. Second, markers of host contamination must be used to evaluate the level of media and host contamination. The number of metabolites with abundances correlated with phenol red or HEPES can be used to assess media contribution. Our studies also suggest that visual detection of ghost material (via microscopy) combined with assessment of host-specific metabolite markers is an effective option to assess sample purity. Finally, data must be normalized to appropriate measurements to maximize metabolome signal associated with the treatment of interest; subsequent subtraction of metabolites associated with host or media (by uninfected erythrocyte control or known media components like HEPES or phenol red) can further reduce metabolite influence by extra-parasite conditions. With these considerations, metabolomics has the potential to be a powerful tool in the study of intracellular parasites.

## MATERIALS AND METHODS

### Parasite Cultivation

Laboratory-adapted *P. falciparum* lines were cultured in RPMI 1640 (Roswell Park Memorial Institute medium, Thermo Fisher Scientific, Waltham, MA) containing HEPES (Sigma Aldrich, St Louis, MO) supplemented with 0.5% AlbuMAX II Lipid-rich BSA (Sigma Aldrich, St Louis, MO) and 50 mg/L hypoxanthine (Thermo Fisher Scientific, Waltham, MA). Parasite cultures were maintained at 3% hematocrit and diluted with human red blood cells (blood batch noted in Table S3) to maintain parasitemia between 1-3%, with change of culture medium every other day (Fig. 1B; **Step 1**). Cultures were incubated at 37°C with 5% oxygen, 5% carbon dioxide and 90% nitrogen (45). Some samples were treated with artemisinin, an antimalarial with metabolic effects (dihydroartemisinin; see Antimalarial Treatment in Table S3) (3, 5).

### Parasite Isolation

Two distinct laboratory-adapted clinical isolates of *P. falciparum* (BEI Resources, NIAID, NIH: *Plasmodium falciparum*, Strain IPC 4884/MRA-1240 and IPC 5202/MRA-1238, contributed by Didier Ménard) containing mixed stages with >50% rings were synchronized using 5% sorbitol (Sigma Aldrich, St Louis, MO) (46). The resultant early stage cultures were incubated at 37°C in AlbuMAX media to allow for the development of a schizont predominant population. After the late stage population was confirmed using microscopy, cultures were checked every one to two hours for the development of newly invaded ring stage parasites. If the parasites were treated with dihydroartemisinin, it was performed at this stage. Fourteen flasks containing early ring-stage parasites (<3 hours post invasion, dihydroartemisinin treated or untreated) were subsequently lysed from the erythrocyte membrane using 0.15% saponin, as previously described (47) (Fig. 1B; **Step 3**). Prior to lysis, sampling of parasite material was taken for determination of erythrocyte count (hemocytometer) and parasitemia (SYBR-green based flow cytometry (48)), which contributed to parasite number determination (total erythrocytes x % parasitemia yields total parasites). Additional samples were obtained following erythrocyte lysis for protein quantification using Bradford reagent (Sigma Aldrich, St Louis, MO). A series of three wash steps were then performed using 1X PBS (Sigma Aldrich, St Louis, MO) using centrifugation at 2000 × g to remove soluble erythrocyte metabolites. Purified material was kept on ice until flash frozen using liquid nitrogen, followed by storage at −80°C until sent for analysis. This procedure was performed five times for both parasite lines (Strain IPC 4884/MRA-1240 and IPC 5202/MRA-1238) to provide 10 drug-treated replicates for metabolomic analysis. Additionally, matched parasites (same parasite lineage, media type, stage, blood batches, and purification methods) were also grown without drug treatment (Table S3) to generate 10 additional control samples (see comparison in Fig. 1A).

### Metabolite Preparation, Analysis, and Identification

Metabolites were identified using Ultrahigh Performance Liquid Chromatography coupled with tandem Mass Spectroscopy (UPLC/MS-MS) by Metabolon, Inc. (Durham, NC). All sample preparations and metabolite identifications were performed according to Metabolon, Inc., standard protocols that are briefly summarized here. Double stranded DNA was quantified in all samples using the Quant-it Picogreen dsDNA Assay Kit (Thermo Fisher, Waltham, MA) according to the manufacturer’s instructions. Recovery standards were added to each sample for extraction quality control purposes. Proteins were precipitated using methanol for 2 min under vigorous shaking then centrifuged for extraction (Fig. 1; **Step 4**). Sample extracts were separated into aliquots, dried, and suspended in appropriate standard-containing solvents for analysis by four methods. These four methods facilitate the measurement of metabolites with different biochemical properties and include reverse phase UPLC/MS-MS methods, one with positive ion electrospray ionization (ESI) optimized for hydrophilic compounds or optimized for hydrophobic compounds and a third method with negative ion mode ESI. Additionally, a UPLC/MS-MS method with negative ion mode ESI following elution from a hydrophilic interaction chromatography column was used. Waters AQUITY ultra-performance liquid chromatography and Thermo Scientific Q-Exactive high resolution/accurate mass spectrometer was used for all metabolite detection (Fig. 1; **Step 5**). Controls that were analyzed in conjunction with the experimental samples included a pooled matrix of all included samples. Internal and recovery standards were used to assess variability and to verify performance of extraction and instrumentation, as routinely performed by Metabolon, Inc.

Raw data was extracted using hardware and software developed by Metabolon, Inc. Metabolites were quantified using area-under-the-curve and identified by comparison to a library of several thousands of pre-existing entries of purified standards or recurrent unknown compounds. Each library standard was uniquely authenticated by retention time/indexes, mass to charge ratios, and chromatographic data. Named metabolites corresponded to library standards or were predicted with confidence according to Metabolon, Inc. standard protocols.

### DNA quantification

Measurement of host derived dsDNA was performed by incubating uninfected erythrocytes at 3% hematocrit for 48 hours in either PBS, RPMI 1640 alone, RPMI 1640 with 50mg/L hypoxanthine, or RPMI with 50mg/L hypoxanthine and 0.5% AlbuMAX II Lipid-rich BSA. Erythrocytes were saponin lysed and washed prior to dsDNA quantification using the Quant-it Picogreen dsDNA Assay Kit as described above.

### Microscopy

Laboratory-adapted *P. falciparum* clones (BEI Resources, NIAID, NIH: *Plasmodium falciparum*, Strain Patient line E/MRA-1000 or IPC 4884/MRA-1238, contributed by Didier Ménard) at 1.5% parasitemia with >60% rings were lysed using 0.15% saponin, as previously described (47). Samples were washed twice using 1X PBS (Sigma Aldrich, St Louis, MO) and centrifugation at 2000 × g for 5 minutes. Samples were then stained on slides with either DAPI at 1:20,000 (Sigma Aldrich, St Louis, MO) and CD235a-PE antibody at 1:100 (Thermo Fisher Scientific, Waltham, MA) for fluorescence microscopy. Fluorescent images were acquired using the EVOS FL Cell Imaging System (Thermo Fisher Scientific, Waltham, MA). Quantification of 1214 parasites associated with erythrocyte membranes was collected in 11 preparations.

### Data Preprocessing and Statistical Analysis

Following the analytical protocol outlined in (49), we first preprocessed metabolite abundances for each sample by imputing missing values with half of the lowest detectable metabolite abundance. Next, we normalized metabolite abundances by sample features (Fig. 3), followed by normalization using metabolite features with log transformation, centering, and scaling (Fig. 1; **Step 6** (50)).

Specifically, to limit inter-sample variability, metabolite abundances for each replicate were normalized to sample value for double stranded DNA, protein, or parasite number. To limit inter-metabolite variability, metabolite abundances were log transformed, centered to median (51), and scaled by standard deviation (Fig 1; **Step 6**).

Resultant processed metabolite abundances were used for univariate and multivariate statistics, as well as classification. All analyses were conducted using R, using tidyverse, knitr, reshape2, pracma, grid, gridExtra, and RSvgDevice for data wrangling and visualization and vegan and base R for analysis (18, 52-63). ANOVAs were used to compare group means for differential abundance determination, and p-values were adjusted using a false discovery rate (Benjamini & Hochberg) to correct for multiple testing (64). The significance cutoff is 0.05. PerMANOVAs were used to compare population separation (Fig 4A-D, F). See supplementary information for code documenting a detailed analysis.

### Random Forest Analysis

Random Forest analysis is a machine learning technique used here to classify sample groups (Fig. 5A). Within a random forest classifier, individual trees are built from subsets of the data, and internally validated on the remaining dataset (Fig. 5A). With this approach, variables (metabolites) are ranked by their effect on classifier accuracy, as measured by a change in performance following removal of the variable (Fig. 5B). Classifiers were built with each data normalization method to predict drug treatment or blood batch. These analyses were conducted in R using the randomForest package and base R (23, 65). See supplementary information for code and detailed analysis.

## ACKNOWLEDGEMENTS

We would like to thank the members of the Guler, Papin, and Petri labs at the University of Virginia for helpful discussion and feedback on experimental design and analysis. Additionally, we would like to thank Michelle Warthan (University of Virginia) for laboratory support.

## SUPPLEMENTAL TABLE/FIGURE LEGENDS

Code available at https://github.com/gulermalaria/metabolomics.

**Table S1:** Metabolomics data. Metabolite abundances provided by Metabolon, Inc., prior to data processing and analysis. See Github specifically (https://github.com/gulermalaria/metabolomics/blob/master/met_methods_raw_data.txt) or Excel spreadsheet.

**Table S2:**
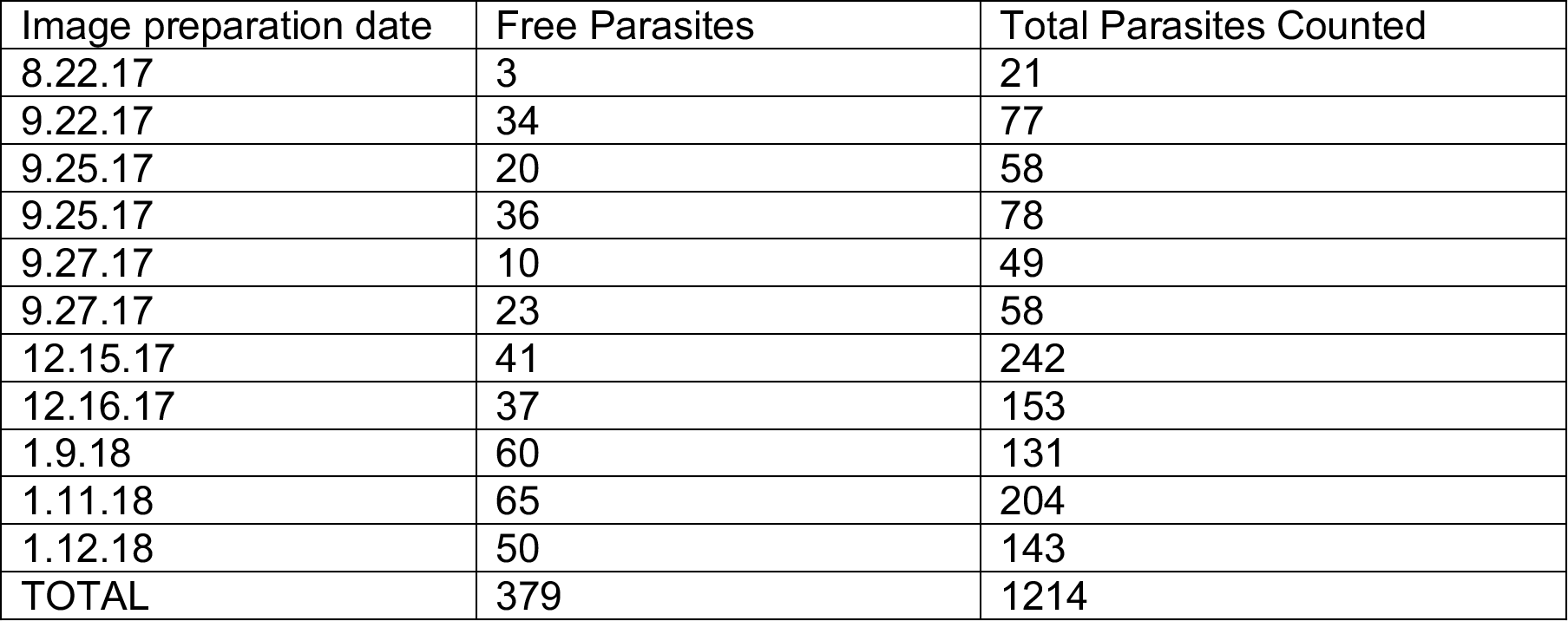
Host entrapment of purified parasites. Parasites remain in host cells following purification. Laboratory adapted *P. falciparum* clones (BEI Resources, NIAID, NIH: *Plasmodium falciparum*, Strain Patient line E/MRA-1000 or IPC 4884/MRA-1238, contributed by Didier Ménard) at >60% rings were lysed using 0.15% saponin, as previously described and in *Materials and Methods.* Samples were washed twice using 1X PBS (Sigma Aldrich, St Louis, MO) and centrifugation at 2000 × g for 5 minutes. Samples were then stained on slides with either DAPI at 1:20,000 (Sigma Aldrich, St Louis, MO) and CD235a-PE antibody at 1:100 (Thermo Fisher Scientific, Waltham, MA) for fluorescence microscopy. Fluorescent images were acquired using the EVOS FL Cell Imaging System (Thermo Fisher Scientific, Waltham, MA). Parasites not associated with erythrocyte membranes were counted.

**Table S3:** Parasite sample reference table. Parasite samples were quantified by protein, DNA, parasite number, parasitemia, and stage distribution.

| Sample ID | Clone | Blood batch | Antimalarial treatment | Protein (mg) | DNA amount (mg/ml) | Parasitemia (%) | Parasite number | Stage distribution |
| --- | --- | --- | --- | --- | --- | --- | --- | --- |
| BAT-A (+) | MRA-1240 | 1 | 700nM DHA | 91.0 | 0.927 | 1.1 | 3.27E+06 | 97% early rings |
| BAT-A (-) | MRA-1240 | 1 | None | 67.1 | 0.587 | 1.1 | 3.27E+06 | 97% early rings |
| BAT-B (+) | MRA-1240 | 1 | 700nM DHA | 121.2 | 0.476 | 1.0 | 2.89E+06 | 98% early rings |
| BAT-B (-) | MRA-1240 | 1 | None | 118.1 | 1.216 | 1.0 | 2.89E+06 | 98% early rings |
| BAT-C (+) | MRA-1240 | 2 | 700nM DHA | 119.2 | 0.985 | 1.0 | 2.93E+06 | 98% early rings |
| BAT-C (-) | MRA-1240 | 2 | None | 87.9 | 1.739 | 1.0 | 2.93E+06 | 98% early rings |
| BAT-D (+) | MRA-1240 | 2 | 700nM DHA | 98.3 | 0.656 | 1.9 | 6.95E+06 | 98% early rings |
| BAT-D (-) | MRA-1240 | 2 | None | 130.1 | 0.557 | 1.9 | 6.95E+06 | 98% early rings |
| BAT-E (+) | MRA-1240 | 3 | 700nM DHA | 125.9 | 1.326 | 2.2 | 6.51E+06 | 93% early rings |
| BAT-E (-) | MRA-1240 | 3 | None | 128.5 | 2.083 | 2.2 | 6.51E+06 | 93% early rings |
| PUR-A (+) | MRA-1238 | 1 | 700nM DHA | 120.2 | 0.325 | 0.6 | 1.31E+06 | 96% early rings |
| PUR-A (-) | MRA-1238 | 1 | None | 125.4 | 0.547 | 0.6 | 1.31E+06 | 96% early rings |
| PUR-B (+) | MRA-1238 | 1 | 700nM DHA | 123.3 | 0.259 | 1.0 | 2.50E+06 | 98% early rings |
| PUR-B (-) | MRA-1238 | 1 | None | 121.0 | 0.673 | 1.0 | 2.50E+06 | 98% early rings |
| PUR-C (+) | MRA-1238 | 2 | 700nM DHA | 104.6 | 0.648 | 0.6 | 2.26E+06 | 97% early rings |
| PUR-C (-) | MRA-1238 | 2 | None | 100.4 | 0.543 | 0.6 | 2.26E+06 | 97% early rings |
| PUR-D (+) | MRA-1238 | 3 | 700nM DHA | 120.7 | 0.599 | 1.0 | 3.32E+06 | 96% early rings |
| PUR-D (-) | MRA-1238 | 3 | None | 110.3 | 1.563 | 1.0 | 3.32E+06 | 96% early rings |
| PUR-E (+) | MRA-1238 | 3 | 700nM DHA | 128.5 | 0.869 | 1.0 | 2.63E+06 | 96% early rings |
| PUR-E (-) | MRA-1238 | 3 | None | 114.5 | 1.118 | 1.0 | 2.63E+06 | 96% early rings |

**Figure S1:**
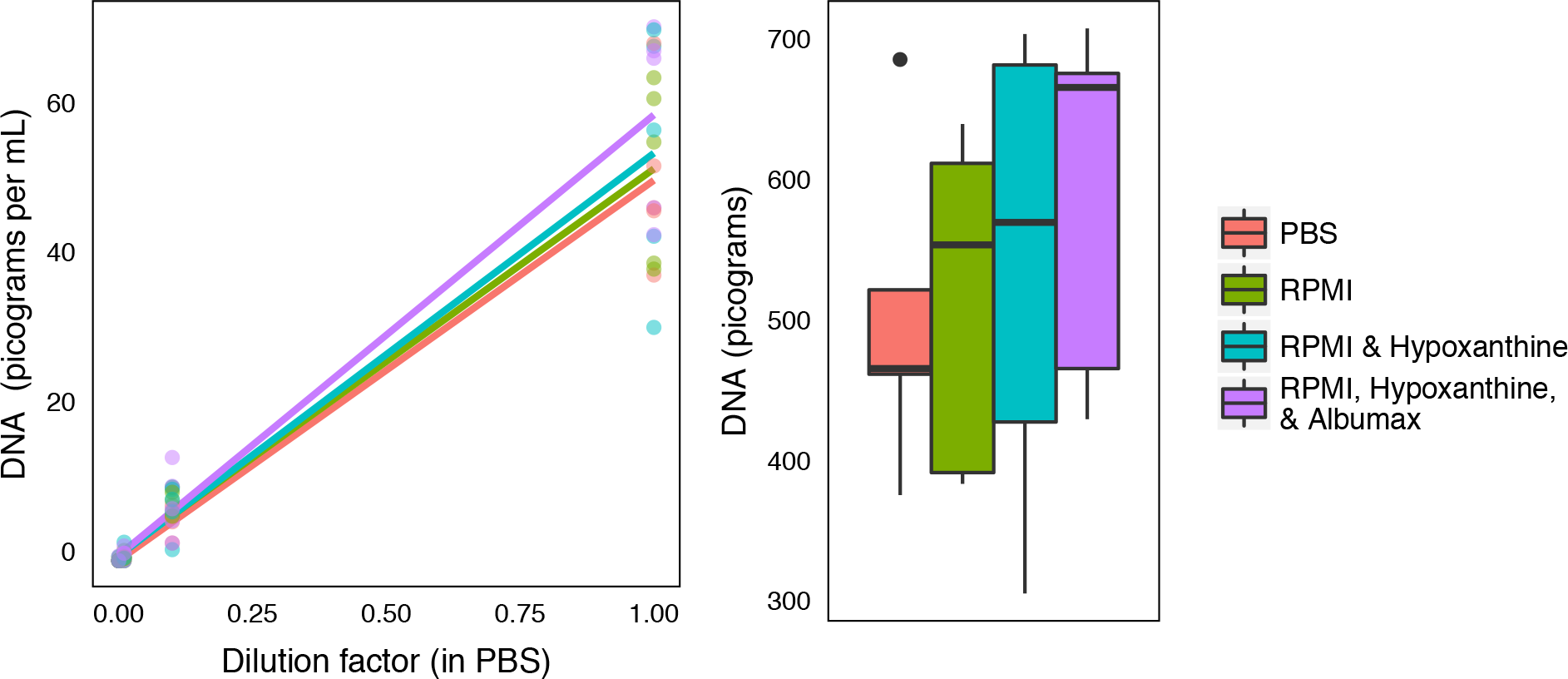
DNA Contribution from host erythrocyte and media. Erythrocytes contribute DNA despite being anucleated. Measurement of host derived dsDNA was performed by incubating uninfected erythrocytes at 3% hematocrit for 48 hours in either PBS, RPMI 1640 alone, RPMI 1640 with 50mg/L hypoxanthine, or RPMI with 50mg/L hypoxanthine and 0.5% AlbuMAX II Lipid-rich BSA. Erythrocytes were saponin lysed and washed twice with PBS prior to dsDNA quantification using the Quant-it Picogreen dsDNA Assay Kit as described in *Materials and Methods.* Nondetectable values (below the limit of detection) were imputed as 0. Left panel demonstrates that DNA abundance is concentration dependent, not mere instrument noise. Right panel demonstrates that media components (like AlbuMAX II Lipid-rich BSA) contribute to DNA quantification, but erythrocytes in PBS contribute the majority of the measured DNA.

